# Virological characteristics of the SARS-CoV-2 KP.3, LB.1 and KP.2.3 variants

**DOI:** 10.1101/2024.06.05.597664

**Authors:** Yu Kaku, Maximilian Stanley Yo, Jarel Elgin Tolentino, Keiya Uriu, Kaho Okumura, The Genotype to Phenotype Japan (G2P-Japan) Consortium, Jumpei Ito, Kei Sato

## Abstract

The SARS-CoV-2 JN.1 variant, arising from BA.2.86.1 with a substitution in the spike (S) protein, S:L455S, exhibited increased fitness and outcompeted the previously predominant XBB lineages by the beginning of 2024. Subsequently, JN.1 subvariants including KP.2 and KP.3, which convergently acquired S protein substitutions such as S:R346T, S:F456L, and S:Q493E, have emerged concurrently. Furthermore, JN.1 subvariants such as LB.1 and KP.2.3, which convergently acquired S:S31del in addition to the above substitutions, have emerged and spread as of June 2024. Here we investigated the virological properties of KP.3, LB.1 and KP.2.3. We estimated the relative effective reproduction number (R_e_) of KP.3, LB.1, and KP.2.3 using a Bayesian multinomial logistic model based on the genome surveillance data from Canada, the UK, and the USA, where these variants have spread from March to April 2024. The R_e_ of KP.3 is more than 1.2-fold higher than that of JN.1 and higher than or comparable to that of KP.2 in these countries. Importantly, the R_e_ values of LB.1 and KP.2.3 are even higher than those of KP.2 and KP.3. These results suggest that the three variants we investigated herein, particularly LB.1, and KP.2.3, will become major circulating variants worldwide in addition to KP.2 and KP.3. The pseudovirus infectivity of KP.2 and KP.3 was significantly lower than that of JN.1. On the other hand, the pseudovirus infectivity of LB.1 and KP.2.3 was comparable to that of JN.1. Neutralization assay was conducted by using four types of breakthrough infection (BTI) sera with XBB.1.5, EG.5, HK.3 and JN.1 infections as well as monovalent XBB.1.5 vaccine sera. In all four groups of BTI sera tested, the 50% neutralization titers (NT50) against LB.1 and KP.2.3 were significantly lower than those against JN.1 (2.2-3.3-fold and 2.0-2.9-fold) and even lower than those against KP.2 (1.6-1.9-fold and 1.4-1.7 fold). Although KP.3 exhibited neutralization resistance against all BTI sera tested than JN.1 (1.6-2.2-fold) with statistical significance, there were no significant differences between KP.3 and KP.2. In the case of infection-naive XBB.1.5 vaccine sera, the NT50 values of JN.1 subvariants were very low. In the case of XBB.1.5 vaccine sera after natural XBB infection, the NT50 values against KP.3, LB.1 and KP.2.3 were significantly lower than those of JN.1 (2.1-2.8-fold) and even lower than KP.2 after infection (1.4-2.0-fold). Overall, our results suggest that the S substitutions convergently acquired in the JN.1 subvariants contribute to immune evasion, and therefore, increase their R_e_ when compared to parental JN.1. More importantly, LB.1 and KP.2.3 exhibited higher pseudovirus infectivity and more robust immune resistance than KP.2. These data suggest that S:S31del is critical to exhibit increased infectivity, increased immune evasion, and therefore, potentially contributes to increased R_e_.

## Text

The SARS-CoV-2 JN.1 variant (BA.2.86.1.1), arising from BA.2.86.1 with a substitution in the spike (S) protein, S:L455S, exhibited increased fitness and outcompeted the previously predominant XBB lineages by the beginning of 2024.^1^ Subsequently, JN.1 subvariants including KP.2 (JN.1.11.1.2) and KP.3 (JN.1.11.1.3), which convergently acquired S protein substitutions such as S:R346T, S:F456L, and S:Q493E, have emerged concurrently (**Figure 1A**).^2^ Furthermore, JN.1 subvariants such as LB.1 (JN.1.9.2.1) and KP.2.3 (JN.1.11.1.2.3), which convergently acquired a deletion at the 31st position in S (S:S31del) in addition to the above substitutions, have emerged and spread as of June 2024 (**Figure 1A**). We have recently reported the virological features of KP.2.^2^ Here we investigated the virological properties of KP.3, LB.1, and KP.2.3.

**Figure 1.**
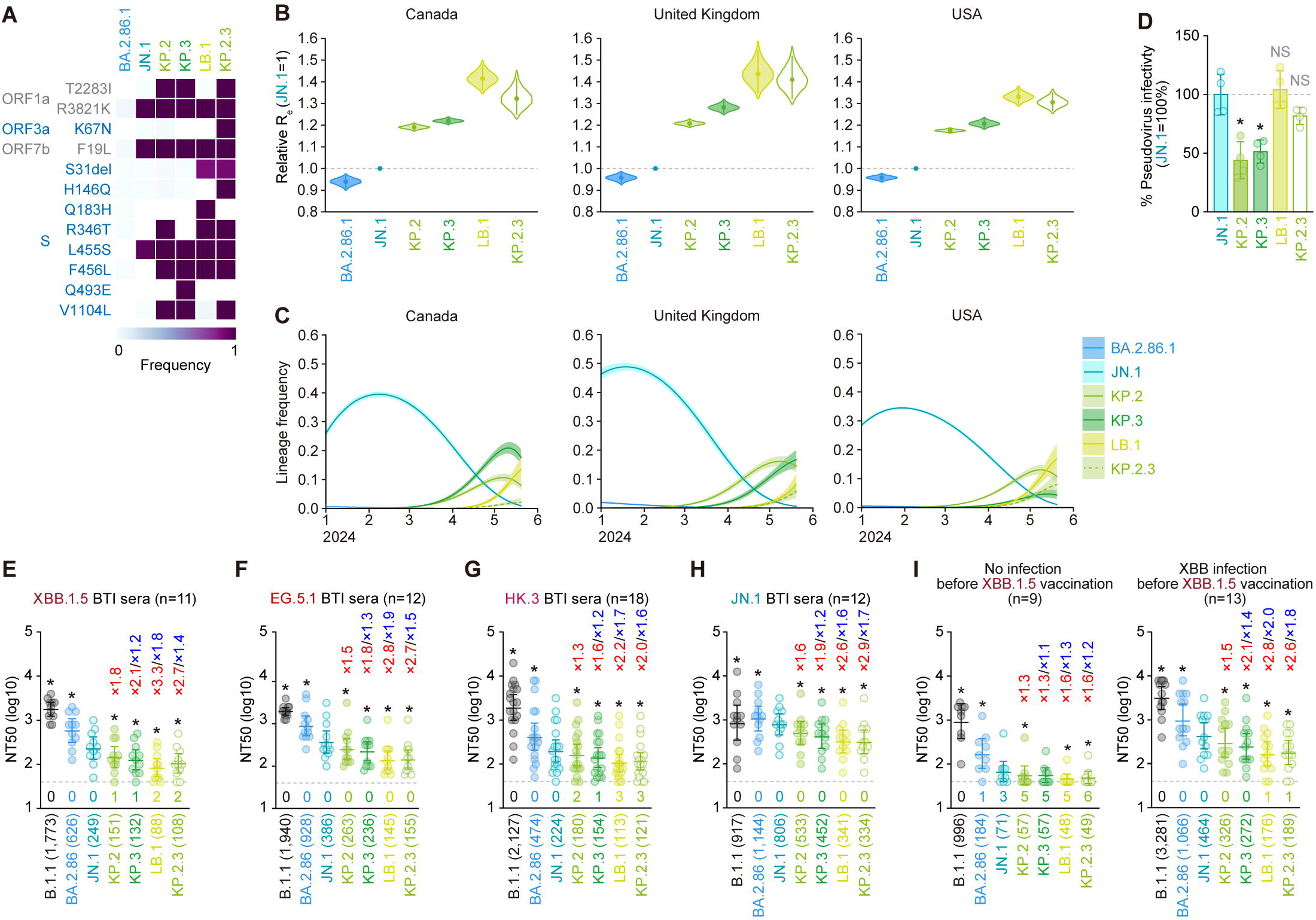
Virological features of KP.3, LB.1 and KP.2.3. (**A**) Frequency of mutations in KP.3, LB.1, KP.2.3, and other lineages of interest. Only mutations with a frequency >0.5 in at least one but not all the representative lineages are shown. (**B**) Estimated relative R_e_ of the variants of interest in Canada, the United Kingdom, and the USA. The relative R_e_ of JN.1 is set to 1 (horizontal dashed line). Violin, posterior distribution; dot, posterior mean; line, 95% credible interval. (**C**) Estimated epidemic dynamics of the variants of interest in Canada, the United Kingdom, and the USA from January 1, 2024 to May 27, 2024. Line, posterior mean, ribbon, 95% credible interval. (**D**) Lentivirus-based pseudovirus assay. HOS-ACE2/TMPRSS2 cells were infected with pseudoviruses bearing each S protein of JN.1, KP.2, KP.3, LB.1 and KP.2.3. The amount of input virus was normalized to the amount of HIV-1 p24 capsid protein. The percentage infectivity of KP.2, KP.3, LB.1 and KP.2.3 are compared to that of JN.1. The horizontal dash line indicates the mean value of the percentage infectivity of JN.1. Assays were performed in quadruplicate, and a representative result of four independent assays is shown. The presented data are expressed as the average ± SD. Each dot indicates the result of an individual replicate. Statistically significant differences versus JN.1 is determined by two-sided Student’s *t* tests and statistically significant differences (*P* < 0.01) versus JN.1 are indicated with asterisks. (**E**-**I**) Neutralization assay. Assays were performed with pseudoviruses harboring the S proteins of B.1.1, BA.2.86, JN.1, KP.2, KP.3, LB.1 and KP.2.3. The following sera were used: convalescent sera from fully vaccinated individuals who had been infected with XBB.1.5 (four 3-dose vaccinated donors, four 4-dose vaccinated donors and three 5-dose vaccinated donors. 11 donors in total) (**E**); EG.5.1 (four 3-dose vaccinated donors, two 4-dose vaccinated donors, two 5-dose vaccinated donors and four 6-dose vaccinated donors. 12 donors in total) (**F**); individuals who had been infected with HK.3 (three 2-dose vaccinated donors, five 3-dose vaccinated donor, two 4-dose vaccinated donors, three 5-dose vaccinated donors, one 6-dose vaccinated donor and four donors with unknown vaccine history. 18 donors in total) (**G**) and individuals who had been infected with JN.1 (one 2-dose vaccinated donor, two 3-dose vaccinated donors, two 7-dose vaccinated donors and seven donors with unknown vaccine history. 12 donors in total) (**H**), and XBB.1.5 monovalent vaccination sera from fully vaccinated individuals who had not been infected (9 donors, **left**) and XBB.1.5 monovalent vaccination sera from fully vaccinated individuals who had been infected with XBB subvariants (after June, 2023) (13 donors, **right**) (**I**). Assays for each serum sample were performed in quadruplicate to determine the 50% neutralization titer (NT_50_). Each dot represents one NT_50_ value, and the geometric mean and 95% confidence interval are shown. The number in parenthesis indicates the geometric mean of NT_50_ values. The horizontal dash line indicates a detection limit (40-fold) and the number of serum donors with the NT_50_ values below the detection limit is shown in the figure (under the bars and dots of each variant). Statistically significant differences (*P* < 0.05) versus JN.1 were determined by two-sided Wilcoxon signed-rank tests and indicated with asterisks. The fold changes of NT_50_ versus JN.1 and KP.2 are indicated with “X” in red and blue.

We estimated the relative effective reproduction number (R_e_) of KP.3, LB.1, and KP.2.3 using a Bayesian multinomial logistic model^3^ based on the genome surveillance data from Canada, the UK, and the USA, where these variants have spread from March to April 2024 (**Figures 1B and 1C; Table S3**). The R_e_ of KP.3 is more than 1.2–fold higher than that of JN.1 and higher than or comparable to that of KP.2 in these countries (**Figure 1B**). Importantly, the R_e_ values of LB.1 and KP.2.3 are even higher than those of KP.2 and KP.3 (**Figure 1B**). These results suggest that the three variants we investigated herein, particularly LB.1, and KP.2.3, will become major circulating variants worldwide in addition to KP.2.

We then performed virological and immunological experiments with pseudoviruses. The pseudovirus infectivity of KP.2 and KP.3 was significantly lower than that of JN.1 (**Figure 1D**). On the other hand, the pseudovirus infectivity of LB.1 and KP.2.3 was comparable to that of JN.1 (**Figure 1D**). Neutralization assay was conducted by using four types of breakthrough infection (BTI) sera with XBB.1.5, EG.5, HK.3 and JN.1 infections as well as monovalent XBB.1.5 vaccine sera. In all four groups of BTI sera tested, the 50% neutralization titers (NT_50_) against LB.1 and KP.2.3 were significantly lower than those against JN.1 (2.2–3.3-fold and 2.0–2.9-fold) and even lower than those against KP.2 (1.6–1.9-fold and 1.4–1.7 fold) (**Figure 1E–1H**). Although KP.3 exhibited neutralization resistance against all BTI sera tested than JN.1 (1.6–2.2-fold) with statistical significance, there were no significant differences between KP.3 and KP.2 (**Figure 1E–1H**). In the case of infection-naïve XBB.1.5 3 vaccine sera, the NT_50_ values of JN.1 subvariants were very low (**Figure 1I**). In the case of XBB.1.5 vaccine sera after natural XBB infection, the NT_50_ values against KP.3, LB.1 and KP.2.3 were significantly lower than those of JN.1 (2.1–2.8-fold) and even lower than KP.2 after infection (1.4–2.0-fold) (**Figure 1I**).

Overall, our results suggest that the S substitutions convergently acquired in the JN.1 subvariants contribute to immune evasion and increased R_e_ compared to the parental JN.1. More importantly, LB.1 and KP.2.3 exhibited higher pseudovirus infectivity and more robust immune resistance than KP.2. These data suggest that S:S31del is critical for increased infectivity and enhanced immune evasion, and potentially contributes to increased R_e_.

## Grants

Supported in part by AMED ASPIRE Program (JP24jf0126002, to G2P-Japan Consortium and Kei Sato); AMED SCARDA Japan Initiative for World-leading Vaccine Research and Development Centers “UTOPIA” (JP243fa627001h0003,to Kei Sato); AMED SCARDA Program on R&D of new generation vaccine including new modality application (JP243fa727002h0003, to Kei Sato); AMED Research Program on Emerging and Re-emerging Infectious Diseases (JP243fa727002, to Kei Sato); JST PRESTO (JPMJPR22R1, to Jumpei Ito);JSPS KAKENHI Fund for the Promotion of Joint International Research (International Leading Research) (JP23K20041, to G2P-Japan Consortium and Kei Sato); JSPS KAKENHI Grant-in-Aid for Early-Career Scientists (JP23K14526, to Jumpei Ito); JSPS KAKENHI Grant-in-Aid for Scientific Research A (JP24H00607, to Kei Sato); Mitsubishi UFJ Financial Group, Inc.Vaccine Development Grant (to Jumpei Ito and Kei Sato); The Cooperative Research Program (Joint Usage/Research Center program) of Institute for Life and Medical Sciences, Kyoto University (to Kei Sato); Japanese Government MEXT Scholarship-Research Category (240042, to Maximilian Stanley Yo;220235, to Jarel Elgin Tolentino).

## References

1. Kaku Y, Okumura K, Padilla-Blanco M, et al. Virological characteristics of the SARS-CoV-2 JN.1 variant. Lancet Infect Dis 2024; 24(2): e82.

2. Kaku Y, Uriu K, Kosugi Y, et al. Virological characteristics of the SARS-CoV-2 KP.2 variant. Lancet Infect Dis 2024.

3. Yamasoba D, Kimura I, Nasser H, et al. Virological characteristics of the SARS-CoV-2 Omicron BA.2 spike. Cell 2022; 185(12): 2103-15.e19.

