## Supplementary Appendix for "Virological characteristics of the SARS-CoV-2 KP.3, LB.1 and KP.2.3 variants"

**Table of Contents**

**Contents** Page


| Contents | Page |
|---|---|
| **Materials and Methods** | 2-5 |
| Ethics statement | |
| Human serum collection | |
| Epidemic dynamics analysis and mutation frequency calculation | |
| Plasmid construction | |
| Cell culture | |
| Pseudovirus preparation | |
| Neutralization assay | |
| Data availability | |
| **Table S1.** Human infection sera used in this study | 6 |
| **Table S2.** Human XBB.1.5 vaccine sera used in this study | 7 |
| **Table S3.** Estimated relative $R_e$ and epidemic dynamics modeling parameters of representative SARS-CoV-2 Omicron sublineages spreading in Canada, the United Kingdom, and the USA from January 1, 2024 to May 27, 2024 | 8-10 |
| **Table S4.** Primers used in this study | 11 |
| **Consortia** | 12 |
| **Acknowledgments** | 13 |
| **Supplemental References** | 14 |

### Materials and Methods

#### Ethics statement

All protocols involving specimens from human subjects recruited at Kyoto University, Interpark Kuramochi Clinic, Namikibashi Clinic, Wakaba Clinic and Keio University were reviewed and approved by the Institutional Review Boards of The Institute of Medical Science, The University of Tokyo (approval IDs: 2021-1-0416 and 2022-29-0915), Kyoto University (approval ID: G1309), Interpark Kuramochi Clinic (approval ID: G2021-004), and Keio University (approval ID: 20200059), respectively. All human subjects provided written informed consent. All protocols for the use of human specimens were reviewed and approved by the Institutional Review Boards of The Institute of Medical Science, The University of Tokyo (approval IDs: 2021-1-0416, 2021-18-0617 and 2022-29-0915).

#### Human serum collection

Convalescent sera were collected from fully vaccinated individuals who had been infected with XBB.1.5 (four 3-dose vaccinated, four 4-dose vaccinated and three 5-dose vaccinated; time interval between the last vaccination and infection, 114–524 days；15–29 days after testing. n=11 in total; average age: 47.1 years, range: 18–74 years, 18.0% male) and EG.5.1 (four 3-dose vaccinated, two 4-dose vaccinated, two 5-dose vaccinated and four 6-dose vaccinated; time interval between the last vaccination and infection, 58–498 days；11–27 days after testing. n=12 in total; average age: 55.7 years, range: 41–77 years, 50% male), individuals who had been infected with HK.3 (three 2-dose vaccinated, five 3-dose vaccinated, two 4-dose vaccinated, three 5-dose vaccinated, one 6-dose vaccinated and four unknown vaccine history; time interval between the last vaccination and infection, 44–888 days；17–112 days after testing. n=18 in total; average age: 57.4 years, range: 10–84 years, 77.8% male) and individuals who had been infected with JN.1 (one 2-dose vaccinated, two 3-dose vaccinated, two 7-dose vaccinated and seven unknown vaccine history; time interval between the last vaccination and infection, 58–498 days；13–46 days after testing. n=12 in total; average age: 69.3 years, range: 31–94 years, 41.7% male). XBB.1.5 monovalent vaccine sera from fully vaccinated individuals who had not been infected (nine donors. Average age, 66.2; range, 51–89; 44.4% male) and those from fully vaccinated individuals who had been infected with XBB subvariants (thirteen donors. Average age, 49.3; range, 32–61; 69.2% male) were collected before vaccination and three-four weeks (20–29 days) after vaccination. The SARS-CoV-2 variants were identified as previously described. [1-4] Sera were inactivated at 56°C for 30 minutes and stored at –80°C until use. The details of the convalescent sera are summarized in **Table S1 and Table S2**.

#### Epidemic dynamics analysis and mutation frequency calculation

In this study, we analyzed the viral genomic surveillance data stored in the GISAID database (https://www.gisaid.org; downloaded on May 29, 2024). We used the data of SARS-CoV-2 collected from January 1, 2024, to May 27, 2024 in this analysis. We excluded any data that i) lacks collection date and PANGO lineage information; ii) was retrieved from non-human animals; iii) was sampled by quarantine; iv) was sampled from the original passage; v) whose genomic sequence is not longer than 28,000 base pairs; and vi) contains >2% of unknown (N) nucleotide sequences. In the downstream analysis, we only used sequences for PANGO lineages with >20 sequences in each country in the dataset. We modeled the epidemic dynamics of variants of interest in Canada, the United Kingdom, and the USA where >100, >50, and >30 genomic sequences of KP.3, LB.1, and KP.2.3 were detected, respectively. The daily frequency of each viral lineage was counted. Then, epidemic dynamics and $R_e$ values for each viral lineage were subsequently estimated according to the Bayesian multinomial logistic model, as described in our previous study. [4] Briefly, we estimated the logistic slope parameter $\beta_l$ for each lineage and then calculated a relative $R_e$ for each lineage ($r_l$) as $r_l = \exp(\gamma\beta_l)$ where $\gamma$ is the average viral generation time (2.1 days) (http://sonorouschocolate.com/covid19/index.php?title=Estimating_Generation_Time_Of_Omicron). For parameter estimation, the intercept and slope parameters of JN.1 were fixed at 0. The relative $R_e$ of JN.1 was fixed at 1, and that of other lineage was estimated with respect to that of JN.1. Parameter estimation was performed by using the Markov chain Monte Carlo (MCMC) approach implemented in CmdStan v2.34.1 (https://mc-stan.org) accessed through the CmdStanr v0.6.1 R interface (https://mc-stan.org/cmdstanr/). Four independent 5,000-step MCMC chains were run including 1,000-step warmup iterations. We confirmed that an estimated $\hat{R}$ convergence diagnostic value is <1.05 and bulk and tail effective sampling sizes are >200, indicating that all runs were successfully convergent. Information on the estimated parameters of all variants is summarized in **Table S3**. Only KP.3, LB.1, and KP.2.3 variants are included in **Figure 1.** Mutation frequency of each lineage was calculated by dividing the number of sequences harboring the substitution of interest with the total number of sequences in each lineage.

#### Plasmid construction

Plasmids expressing the SARS-CoV-2 spike proteins of B.1.1, BA.2.86, JN.1 and KP.2 were prepared in our previous studies. [5-8] Plasmids expressing the spike protein of KP.2.3, KP.3 and LB.1 were generated by site-directed overlap extension PCR using pC-SARS2-S JN.1 and KP.2 as the template and the primers listed in **Table S4**. The resulting PCR fragment was subcloned into the KpnI-NotI site of the pCAGGS vector[9] using In-Fusion HD Cloning Kit (Takara, Cat# Z9650N). Nucleotide sequences were determined by DNA sequencing services (Eurofins), and the sequence data were analyzed by SnapGene software v6.1.1 (www.snapgene.com).

#### Cell culture

The Lenti-X 293T cells (Takara, Cat# 632180) and HOS-ACE2/TMPRSS2 cells (kindly provided by Dr. Kenzo Tokunaga), a derivative of HOS cells (a human osteosarcoma cell line; ATCC CRL-1543) stably expressing human ACE2 and TMPRSS2,[10,11] were maintained in Dulbecco's modified Eagle's medium (DMEM) (high glucose) (Wako, Cat# 044- 29765) containing 10% fetal bovine serum (Sigma-Aldrich Cat# 172012-500ML), 100 units penicillin and 100 ug/ml streptomycin (Sigma-Aldrich, Cat# P4333-100ML).

**Pseudovirus preparation**

Pseudoviruses were prepared as previously described. [5,7,8,12] Briefly, lentivirus (HIV-1)-based, luciferase-expressing reporter viruses were pseudotyped with the SARS-CoV-2 S. One prior day of transfection, the LentiX-293T cells were seeded at a density of 2 × $10^6$ cells. The LentiX-293T cells were cotransfected with 1 μg psPAX2-IN/HiBiT (a packaging plasmid encoding the HiBiT-tag-fused integrase[10]), 1 μg pWPI-Luc2 (a reporter plasmid encoding a firefly luciferase gene[13]) and 500 ng plasmids expressing parental S or its derivatives using TransIT-293 transfection reagent (Mirus, Cat# MIR2704) according to the manufacturer's protocol. Two days post transfection, the culture supernatants were harvested and filtrated. The amount of produced pseudovirus particles was quantified by the HiBiT assay using Nano Glo HiBiT lytic detection system (Promega, Cat# N3040) as previously described. [13] In this system, HiBiT peptide is produced with HIV-1 integrase and forms NanoLuc luciferase with LgBiT, which is supplemented with substrates. In each pseudovirus particle, the detected HiBiT value is correlated with the amount of the pseudovirus capsid protein, HIV-1 p24 protein. [13] Therefore, we calculated the amount of HIV-1 p24 capsid protein based on the HiBiT value measured, according to the previous paper. [13] To measure viral infectivity, the same amount of pseudovirus normalized with the HIV-1 p24 capsid protein was inoculated into HOS-ACE2/TMPRSS2 cells. At two days postinfection, the infected cells were lysed with a Bright-Glo luciferase assay system (Promega, Cat# E2620), and the luminescent signal produced by firefly luciferase reaction was measured using a GloMax explorer multimode microplate reader 3500 (Promega). The pseudoviruses were stored at –80°C until use.

**Neutralization assay**

Neutralization assays were performed previously described[5-8] with some modifications. First, the assays were mainly conducted by a semi-automated high-throughput method using Fluent780 (Tecan). [14,15] The SARS-CoV-2 spike pseudoviruses (counting ~100,000 relative light units) and serially diluted (40-fold to 29,160-fold dilution at the final concentration) heat-inactivated sera were manually prepared in a 2-ml 96-well plate (Greiner, Cat# 780271) and in 96-well microplates (ThermoFisher Scientific, Cat# 168136), respectively. The pseudoviruses were dispensed and mixed with the sera in 384-well plates (ThermoFisher Scientific, Cat# 164610) on Fluent780 (Tecan). Pseudoviruses without sera were included as controls. After incubation at 37°C for 1 hour, HOS-ACE2/TMPRSS2 cells (3,000 cells/30 μl) were added to the 20 μl mixture of pseudovirus and serum in the 384-well white plate on the device. Two days post infection, the infected cells were lysed with a Bright-Glo luciferase assay system (Promega, Cat# E2620) on Fluent780 (Tecan), and the luminescent signal was measured and processed using an Infinite200 and a Magellan (Tecan). The assay of each serum sample was performed in quadruplicate, and the 50% neutralization titer ($NT_{50}$) was calculated using Prism 9 (GraphPad Software).

**Data availability**

The GISAID datasets used in this study are available from the GISAID database (https://www.gisaid.org; EPI-SET-ID: EPI_SET_240529sa). The supplemental tables for the GISAID datasets are available in the GitHub repository (https://github.com/TheSatoLab/KP.3_short).

**Table S1. Human infection sera used in this study**

| SARS-CoV-2 infected | Donor ID | Sex | Age | Date of 1st vaccination (YYYY-MM-DD) | Date of 2nd vaccination (YYYY-MM-DD) | Date of 3rd vaccination (YYYY-MM-DD) | Date of 4th vaccination (YYYY-MM-DD) | Date of 5th vaccination (YYYY-MM-DD) | Date of 6th vaccination (YYYY-MM-DD) | Date of 7th vaccination (YYYY-MM-DD) | Date of test (YYYY-MM-DD) | Date of sampling (YYYY-MM-DD) | Prior infection? |
|---|---|---|---|---|---|---|---|---|---|---|---|---|---|
| XBB.1.5 | 37306 | Female | 53 | 2021-04-27 (P) | 2021-05-18 (P) | 2022-02-01 (P) | 2022-07-30 (M) | 2022-12-17 (P) | | | 2023-07-20 | 2023-08-08 | No |
| XBB.1.5 | 37097 | Female | 44 | NA (M) | 2021-08-16 (M) | 2022-05-13 (M) | | | | | 2023-07-13 | 2023-08-11 | No |
| XBB.1.5 | 37071 | Female | 48 | 2021-10-01 (P) | 2021-11-01 (P) | 2022-05-06 (P) | | | | | 2023-07-15 | 2023-08-11 | No |
| XBB.1.5 | 37229 | Female | 74 | 2021-06-24 (P) | 2021-07-15 (P) | 2022-02-16 (M) | 2022-07-20 (M) | 2023-03-25 (M) | | | 2023-07-17 | 2023-08-01 | No |
| XBB.1.5 | 38084 | Male | 44 | 2021-09-13 (P) | 2021-10-05 (P) | 2022-07-29 (M) | | | | | 2023-08-11 | 2023-09-02 | No |
| XBB.1.5 | 38019 | Female | 18 | 2021-09-07 (P) | 2021-10-07 (P) | 2022-04-28 (P) | 2022-12-27 (P) | | | | 2023-08-10 | 2023-09-04 | No |
| XBB.1.5 | 38952 | Female | 54 | 2021-07-27 (M) | 2021-08-24 (M) | 2022-03-24 (P) | 2022-10-26 (P) | | | | 2023-08-23 | 2023-09-10 | No |
| XBB.1.5 | 38880 | Male | 51 | 2021-10-07 (P) | 2021-10-28 (P) | 2022-05-13 (M) | 2022-11-09 (PBA4/5) | | | | 2023-08-22 | 2023-09-16 | No |
| XBB.1.5 | 39019 | Female | 22 | 2021-07-26 (M) | 2021-08-23 (M) | 2022-03-19 (P) | | | | | 2023-08-25 | 2023-09-13 | No |
| XBB.1.5 | 39502 | Female | 43 | 2021-07-31 (P) | 2021-08-23 (P) | 2022-03-15 (P) | 2022-11-01 (P) | | | | 2023-09-05 | 2023-09-26 | No |
| XBB.1.5 | 39321 | Female | 67 | 2021-06-14 (P) | 2021-07-05 (P) | 2022-02-10 (M) | 2022-07-26 (M) | 2023-01-06 (P) | | | 2023-09-01 | 2023-09-27 | No |
| EG.5 | 38111 | Female | 50 | 2021-08-16 (P) | 2021-09-06 (P) | 2022-04-01 (P) | | | | | 2023-08-12 | 2023-09-02 | No |
| EG.5 | 38197 | Female | 49 | 2021-08-20 (P) | 2021-09-10 (P) | 2022-03-25 (M) | 2022-11-18 (P) | | | | 2023-08-13 | 2023-09-02 | No |
| EG.5 | 38217 | Male | 62 | 2021-07-04 (P) | 2021-07-25 (P) | 2022-02-26 (M) | 2022-08-06 (M) | 2022-11-20 (P) | | | 2023-08-13 | 2023-09-02 | No |
| EG.5 | 37999 | Male | 67 | 2021-06-07 (P) | 2021-07-01 (P) | 2022-02-08 (P) | 2022-07-12 (P) | 2022-12-24 (PBA4/5) | 2023-06-13 (MBA4/5) | | 2023-08-10 | 2023-09-02 | No |
| EG.5 | 39025 | Male | 63 | 2021-08-01 (P) | 2021-08-22 (P) | 2022-03-07 (M) | 2022-08-10 (M) | 2022-11-26 (PBA4/5) | 2023-06-06 (PBA4/5) | | 2023-08-26 | 2023-09-11 | No |
| EG.5 | 39288 | Male | 77 | 2021-06-09 (P) | 2021-07-01 (P) | 2022-02-05 (M) | 2022-07-12 (P) | 2022-11-18 (PBA4/5) | 2023-05-30 (PBA1) | | 2023-08-31 | 2023-09-20 | No |
| EG.5 | 39301 | Male | 41 | 2021-08-26 (M) | 2021-09-23 (M) | 2022-04-24 (M) | 2022-10-09 (MBA1) | | | | 2023-09-01 | 2023-09-23 | No |
| EG.5 | 39314 | Female | 56 | 2021-07-25 (P) | 2021-08-22 (P) | 2022-03-03 (M) | 2022-08-06 (M) | 2022-11-06 (M) | 2023-04-01 (NA) | | 2023-09-01 | 2023-09-21 | No |
| EG.5 | 39315 | Female | 60 | 2021-08-29 (P) | 2021-09-19 (P) | 2022-04-16 (M) | 2022-09-16 (P) | 2022-12-16 (PBA4/5) | | | 2023-08-31 | 2023-09-23 | No |
| EG.5 | 39328 | Female | 59 | 2021-09-30 (P) | 2021-10-27 (P) | 2022-05-17 (P) | | | | | 2023-08-31 | 2023-09-27 | Yes |
| EG.5 | KK | Male | 43 | 2021-06-17 (P) | 2021-07-08 (P) | 2022-09-02 (P) | | | | | 2023-09-22 | 2023/10/03 | No |
| EG.5 | RK | Female | 41 | 2021-06-16 (P) | 2021-07-07 (P) | 2022-09-02 (P) | | | | | 2023-09-22 | 2023/10/03 | No |
| HK.3 | 42405 | Male | 56 | NA | 2021-08-27 (P) | 2022-03-14 (M) | 2022-08-22 (P) | 2022-12-01 (P) | 2023-07-26 (P) | | 2023-11-03 | 2024-02-17 | Yes |
| HK.3 | 42410 | Male | 66 | 2021-04-23 (P) | 2021-05-14 (P) | 2022-01-27 (P) | 2022-08-10 (P) | 2022-12-07 (M) | | | 2023-11-04 | 2024-02-13 | No |
| HK.3 | 42412 | Female | 75 | 2021-05-27 (P) | 2021-06-17 (P) | 2022-01-29 (P) | 2022-12-16 (P) | 2023-05-26 (P) | | | 2023-11-05 | 2024-02-18 | No |
| HK.3 | 42413 | Male | 10 | 2022-03-25 (P) | 2022-07-22 (P) | 2023-01-17 (P) | | | | | 2023-11-05 | 2024-02-18 | No |
| HK.3 | 42414 | Female | 41 | 2021-10-29 (P) | 2021-11-22 (P) | 2022-05-28 (P) | 2022-11-08 (P) | 2023-12-10 (P) | | | 2023-11-05 | 2024-02-18 | No |
| HK.3 | 42418 | Female | 25 | 2021-03-18 (P) | 2021-04-07 (P) | 2022-01-20 (P) | | | | | 2023-11-05 | 2024-02-16 | No |
| HK.3 | 42423 | Male | 49 | 2021-07-06 (M) | 2021-08-10 (M) | 2022-03-12 (M) | 2022-09-20 (M) | | | | 2023-11-05 | 2024-02-25 | No |
| HK.3 | 42429 | Male | 66 | 2021-07-05 (P) | 2021-07-26 (P) | 2022-03-04 (P) | | | | | 2023-11-06 | 2024-02-15 | NA |
| HK.3 | 39335 | Male | 50 | 2021-09-09 (P) | 2021-09-30 (P) | 2022-05-09 (P) | 2022-12-16 (P) | | | | 2023-08-31 | 2023-09-23 | NA |
| HK.3 | 39339-1 | Male | 53 | 2021-10-01 (P) | 2022-06-01 (P) | | | | | | 2023-09-02 | 2023-09-23 | NA |
| HK.3 | 3319 | Female | 55 | 2021-10-01 (P) | 2021-10-22 | 2022-07-10 | | | | | 2023-12-13 | 2024-01-17 | NA |
| HK.3 | 3363 | Male | 38 | NA | 2021-08 | | | | | | 2024-01-06 | 2024-01-27 | NA |
| HK.3 | 3373 | Male | 73 | 2022 | 2022 | | | | | | 2024-01-11 | 2024-02-01 | No |
| HK.3 | 3378 | Male | 75 | NA | 2021-06-11 | NA | | | | | 2024-01-14 | 2024-02-07 | NA |
| HK.3 | 3342 | Male | 76 | - | | | | | | | 2023-12-29 | 2024-01-19 | NA |
| HK.3 | 3353 | Male | 83 | - | | | | | | | 2024-01-02 | 2024-01-26 | NA |
| HK.3 | 3354 | Male | 84 | - | | | | | | | 2024-01-02 | 2024-01-19 | NA |
| HK.3 | 3391 | Male | 58 | - | | | | | | | 2024-01-20 | 2024-02-15 | NA |
| JN.1 | 3315 | Female | 75 | NA | NA | NA | NA | NA | NA | 2023-11-07 | 2023-12-11 | 2024-01-15 | No |
| JN.1 | 3323 | Female | 73 | 2021 | 2021 | 2021-05 | | | | | 2023-12-15 | 2023-12-28 | Yes |
| JN.1 | 3325 | Female | 73 | 2021-06 | 2021-07 | | | | | | 2023-12-16 | 2024-01-16 | NA |
| JN.1 | 3338 | Male | 31 | 2021-06 | 2021-07 | 2022-02-02 | | | | | 2023-12-26 | 2024-02-10 | Yes |
| JN.1 | 3355 | Male | 54 | NA | NA | NA | NA | NA | NA | NA | 2024-01-03 | 2024-02-15 | NA |
| JN.1 | 3316 | Female | 52 | - | | | | | | | 2023-12-11 | 2024-01-06 | NA |
| JN.1 | 3320 | Female | 80 | - | | | | | | | 2023-12-13 | 2024-01-12 | NA |
| JN.1 | 3329 | Male | 94 | - | | | | | | | 2023-12-18 | 2024-01-23 | NA |
| JN.1 | 3337 | Male | 72 | - | | | | | | | 2023-12-26 | 2024-01-19 | NA |
| JN.1 | 3362 | Male | 74 | - | | | | | | | 2024-01-06 | 2024-01-19 | NA |
| JN.1 | 3367 | Female | 70 | - | | | | | | | 2024-01-08 | 2024-01-26 | NA |
| JN.1 | 3400 | Female | 84 | - | | | | | | | 2024-01-24 | 2024-02-09 | NA |

NA, not applicable.

P, Pfizer-BioNTech; M, Moderna

**Table S2. Human XBB.1.5 vaccine sera used in this study**

| Donor ID | Sex | Age | Date of 1st vaccination (YYYY-MM-DD) | Date of 2nd vaccination (YYYY-MM-DD) | Date of 3rd vaccination (YYYY-MM-DD) | Date of 4th vaccination (YYYY-MM-DD) | Date of 5th vaccination (YYYY-MM-DD) | Date of 6th vaccination (YYYY-MM-DD) | Date of sampling (before vaccination) (YYYY-MM-DD) | Date of XBB.1.5 vaccination (YYYY-MM-DD) | Date of sampling (after vaccination) (YYYY-MM-DD) | Time interval between vaccination and the second sampling | Prior infection? | variant |
|---|---|---|---|---|---|---|---|---|---|---|---|---|---|---|
| 5165 | Female | 89 | 2021-05-29 (P) | 2021-06-21 (P) | 2022-02-16 (P) | 2022-07-17 (P) | 2022-11-27 (PBA4/5) | 2023-05-20 (PBA4/5) | 2023-09-29 | 2023-09-29 (XBB1.5) | 2023-10-26 | 27 | No | |
| 5166 | Male | 77 | 2021-06-05 (P) | 2021-07-04 (P) | 2022-03-05 (P) | 2022-08-06 (P) | 2022-11-20 (PBA4/5) | 2023-05-20 (PBA4/5) | 2023-09-29 | 2023-09-29 (XBB1.5) | 2023-10-21 | 22 | No | |
| 6783 | Male | 57 | 2021-06-23 (M) | 2021-07-21 (M) | 2022-02-11 (M) | 2022-10-15 (MBA1) | | | 2023-10-03 | 2023-10-03 (XBB1.5) | 2023-10-23 | 20 | No | |
| 2477 | Male | 81 | 2021-07-10 (P) | 2021-07-31 (P) | 2022-03-06 (P) | 2022-08-06 (P) | 2022-11-07 (PBA4/5) | 2023-05-09 (PBA4/5) | 2023-09-29 | 2023-09-29 (XBB1.5) | 2023-10-28 | 29 | No | |
| 6858 | Female | 62 | 2021-07-18 (P) | 2021-08-11 (P) | 2022-02-27 (M) | 2022-07-30 (P) | 2022-11-20 (PBA4/5) | | 2023-10-07 | 2023-10-07 (XBB1.5) | 2023-10-28 | 21 | No | |
| 192 | Female | 64 | 2021-04-22 (P) | 2021-05-13 (P) | 2022-01-15 (P) | 2022-07-16 (P) | 2022-11-24 (PBA4/5) | 2023-05-25 (PBA4/5) | 2023-10-02 | 2023-10-26 (PXBB1.5) | 2023-11-20 | 25 | No | |
| 1700 | Female | 53 | 2021-04-21 (P) | 2021-05-12 (P) | 2022-01-15 (P) | 2022-07-13 (P) | 2022-11-30 (PBA4/5) | 2023-06-23 (MBA4/5) | 2023-09-29 | 2023-10-18 (PXBB1.5) | 2023-11-13 | 26 | No | |
| 5555 | Female | 51 | 2021-07-14 (P) | 2021-08-14 (P) | 2022-02-22 (P) | 2022-07-23 (P) | 2022-12-03 (PBA4/5) | 2023-05-11 (PBA4/5) | 2023-09-30 | 2023-10-21 (PXBB1.5) | 2023-11-15 | 25 | No | |
| 5986 | Male | 62 | 2021-07-24 (P) | 2021-08-14 (P) | 2022-03-12 (M) | 2022-08-27 (M) | 2022-12-24 (PBA4/5) | 2023-06-03 (PBA4/5) | 2023-09-25 | 2023-10-21 (PXBB1.5) | 2023-11-13 | 23 | No | |
| KS | Male | 41 | 2021-06-17 (P) | 2021-07-07 (P) | 2022-03-28 (M) | 2022-10-27 (MBA.5) | | | 2023-09-19 | 2023-09-20 (XBB1.5) | 2023-10-14 | 24 | Yes (2023-06-29) | XBB.1.9 |
| KY | Female | 53 | 2021-08-18 (P) | 2021-09-08 (P) | 2022-04-13 (P) | 2022-10-21 (P) | | | 2023-09-25 | 2023-09-27 (XBB1.5) | 2023-10-20 | 23 | Yes (2023-07-24) | XBB.1.16 |
| KK | Male | 56 | 2021-07-16 (P) | 2021-08-06 (P) | 2022-03-03 (M) | 2022-08-09 (P) | | | 2023-09-25 | 2023-09-29 (XBB1.5) | 2023-10-24 | 25 | Yes (2023-07-17) | EG.5 |
| 2345 | Female | 52 | 2021-07-11 (P) | 2021-08-01 (P) | 2022-03-10 (P) | 2022-10-22 (PBA1) | | | 2023-10-06 | 2023-10-06 (XBB1.5) | 2023-10-27 | 21 | Yes (2023-07) | NA |
| 80 | Male | 61 | 2021-05-10 (P) | 2021-05-31 (P) | 2022-01-24 (M) | 2022-07-22 (BA4/5) | 2022-11-12 (BA4/5) | 2023-06-03 (BA4/5) | 2023-09-29 | 2023-09-30 (XBB1.5) | 2023-10-24 | 24 | Yes (2023-07) | NA |
| 90 | Male | 47 | 2021-05-11 (P) | 2021-06-02 (P) | 2022-01-25 (M) | 2022-08-02 (BA4/5) | 2022-11-09 (BA4/5) | | 2023-10-10 | 2023-10-11 (XBB1.5) | 2023-11-01 | 21 | Yes (2023-07) | NA |
| 100 | Male | 32 | 2021-05-18 (P) | 2021-06-08 (P) | 2022-01-31 (M) | 2022-07-17 (MBA4/5) | 2022-11-15 (PBA4/5) | 2023-05-27 (MBA4/5) | 2023-10-11 | 2023-10-14 (XBB1.5) | 2023-11-06 | 23 | Yes (2023-06) | NA |
| 286691 | Male | 49 | 2021-06-01(P) | 2021-06-22 (P) | 2022-03-21 (M) | | | | 2023-10-19 | 2023-10-19 (PXBB1.5) | 2023-11-16 | 28 | Yes (2023-08-22) | XBB.1.9 |
| 2439306 | Male | 43 | 2021-03-08 (P) | 2021-03-29 (P) | 2021-12-20 (P) | 2022-08-26 (M) | 2022-11-30 (M) | | 2023-10-23 | 2023-10-23 (PXBB1.5) | 2023-11-20 | 28 | Yes (2023-09-08) | EG.5 |
| 4177 | Female | 49 | 2021-04-24 (P) | 2021-05-15 (P) | 2022-01-22 (P) | 2022-07-09 (P) | 2022-12-03 (PBA4/5) | 2023-06-13 (MBA4/5) | 2023-09-29 | 2023-10-14 (PXBB1.5) | 2023-11-10 | 27 | Yes (2023-08-29) | NA |
| 38579 | Male | 48 | 2022-05-28 (NA) | | | | | | 2023-10-31 | 2023-10-31 (XBB1.5) | 2023-11-26 | 26 | Yes (2023-08-16) | XBB.1.16 |
| 36708 | Male | 55 | 2021-08-07 (P) | 2021-08-27 (P) | 2022-04-14 (P) | | | | 2023-10-21 | 2023-11-11 (XBB1.5) | 2023-12-09 | 28 | Yes (2023-05-31) | XBB.1.5 |
| 38061 | Female | 55 | 2021-08-17 (P) | 2021-09-18 (P) | 2022-04-02 (M) | 2022-10-14 (PBA1) | | | 2023-11-28 | 2023-11-28 (XBB1.5) | 2023-12-19 | 21 | Yes (2023-08-10) | XBB.1.5 |

NA, not applicable.

P, Pfizer-BioNTech; M, Moderna

**Table S3. Estimated relative Re and epidemic dynamics modeling parameters of representative SARS-CoV-2 Omicron sublineages spreading in Canada, the United Kingdom, and the USA from January 1, 2024 to May 27, 2024**

| PANGO lineage | Country | Relative $R_e$ (posterior values) Mean | 2.5[th] percentile | 97.5[th] percentile | $\hat{R}$ | Bulk effective sample size | Tail effective sample size |
|---|---|---|---|---|---|---|---|
| KP.3.2 | Canada | 1.719 | 1.581 | 1.877 | 1.000 | 38148.8 | 10912.5 |
| KP.3.1 | Canada | 1.705 | 1.497 | 1.960 | 1.000 | 39667.9 | 11615.7 |
| LB.1 | Canada | 1.416 | 1.360 | 1.478 | 1.001 | 39149.8 | 11075.5 |
| XDV.1 | Canada | 1.385 | 1.283 | 1.510 | 1.001 | 38003.9 | 11057.4 |
| KP.2.3 | Canada | 1.325 | 1.247 | 1.419 | 1.001 | 37842.3 | 11378.9 |
| KP.1.1.1 | Canada | 1.266 | 1.226 | 1.310 | 1.000 | 37403.1 | 12083.4 |
| JN.1.16.1 | Canada | 1.247 | 1.208 | 1.290 | 1.001 | 36824.7 | 12403.0 |
| KP.3 | Canada | 1.220 | 1.206 | 1.234 | 1.000 | 29301.5 | 11177.5 |
| KS.1 | Canada | 1.215 | 1.190 | 1.241 | 1.000 | 33927.6 | 11309.4 |
| KP.2 | Canada | 1.191 | 1.177 | 1.206 | 1.000 | 31135.1 | 12393.9 |
| KP.2.1 | Canada | 1.190 | 1.150 | 1.236 | 1.000 | 40800.9 | 10209.5 |
| JN.1.26 | Canada | 1.189 | 1.153 | 1.231 | 1.000 | 35683.9 | 11317.8 |
| KP.4.1 | Canada | 1.188 | 1.147 | 1.233 | 1.000 | 33958.7 | 11083.6 |
| JN.1.1.6 | Canada | 1.175 | 1.141 | 1.213 | 1.000 | 39058.1 | 11618.0 |
| KP.1.1 | Canada | 1.174 | 1.154 | 1.195 | 1.000 | 35798.1 | 11574.0 |
| JN.1.16 | Canada | 1.159 | 1.142 | 1.178 | 1.000 | 34401.4 | 10925.0 |
| XDK.1 | Canada | 1.146 | 1.125 | 1.170 | 1.002 | 38051.1 | 11346.3 |
| JN.1.11.1 | Canada | 1.133 | 1.119 | 1.148 | 1.002 | 32765.4 | 11486.9 |
| JN.1.32 | Canada | 1.107 | 1.095 | 1.119 | 1.000 | 36420.7 | 12173.6 |
| XDK | Canada | 1.104 | 1.085 | 1.124 | 1.001 | 35905.9 | 10355.5 |
| JN.1.13.1 | Canada | 1.101 | 1.091 | 1.112 | 1.000 | 32736.2 | 12226.4 |
| JN.1.7 | Canada | 1.092 | 1.086 | 1.099 | 1.000 | 26002.3 | 12597.2 |
| XDP | Canada | 1.090 | 1.079 | 1.102 | 1.000 | 32970.5 | 11977.8 |
| JN.1.7.2 | Canada | 1.087 | 1.073 | 1.101 | 1.000 | 35643.4 | 12505.6 |
| JN.1.4.4 | Canada | 1.081 | 1.066 | 1.097 | 1.000 | 32960.1 | 11620.5 |
| JN.1.18 | Canada | 1.076 | 1.063 | 1.089 | 1.000 | 36766.6 | 11900.0 |
| JN.1.20 | Canada | 1.075 | 1.053 | 1.097 | 1.001 | 36248.9 | 11676.6 |
| JN.1.4.6 | Canada | 1.069 | 1.058 | 1.081 | 1.000 | 32524.1 | 12045.9 |
| JN.1.34 | Canada | 1.056 | 1.038 | 1.074 | 1.000 | 35348.9 | 11810.7 |
| JN.1.8.1 | Canada | 1.054 | 1.048 | 1.060 | 1.000 | 27806.2 | 12767.2 |
| JN.1.49 | Canada | 1.042 | 1.016 | 1.067 | 1.000 | 41100.6 | 11046.1 |
| KP.1 | Canada | 1.039 | 1.018 | 1.060 | 1.001 | 40225.5 | 11908.6 |
| JN.1.4 | Canada | 1.033 | 1.029 | 1.037 | 1.000 | 23401.0 | 13604.9 |
| JN.1.46 | Canada | 1.028 | 1.007 | 1.048 | 1.000 | 34294.7 | 12144.0 |
| JN.1.4.2 | Canada | 1.023 | 0.996 | 1.049 | 1.001 | 34495.6 | 11761.0 |
| JN.1.19 | Canada | 1.022 | 0.999 | 1.045 | 1.000 | 35936.0 | 10343.5 |
| JN.1.15 | Canada | 1.022 | 0.996 | 1.048 | 1.000 | 39226.4 | 12113.6 |
| JN.1.8 | Canada | 1.021 | 1.006 | 1.035 | 1.000 | 35261.3 | 11535.9 |
| JN.1.39 | Canada | 1.017 | 1.006 | 1.027 | 1.000 | 35494.5 | 11846.2 |
| JN.1.4.5 | Canada | 1.013 | 1.007 | 1.020 | 1.000 | 31355.2 | 12182.0 |
| JN.1.4.7 | Canada | 1.012 | 0.992 | 1.031 | 1.000 | 36704.0 | 11888.2 |
| JN.1.38 | Canada | 1.011 | 0.984 | 1.037 | 1.000 | 37229.6 | 11686.9 |
| JN.1.5 | Canada | 1.006 | 0.990 | 1.023 | 1.000 | 36437.1 | 11995.0 |
| JN.1.9 | Canada | 1.005 | 0.993 | 1.017 | 1.000 | 34322.2 | 11602.6 |
| JN.1.31 | Canada | 0.997 | 0.989 | 1.005 | 1.000 | 33747.4 | 11268.6 |
| JN.1.11 | Canada | 0.993 | 0.958 | 1.026 | 1.001 | 36204.4 | 11073.6 |
| JN.1.42 | Canada | 0.985 | 0.966 | 1.004 | 1.000 | 32365.0 | 11334.1 |
| JN.1.3 | Canada | 0.985 | 0.954 | 1.015 | 1.000 | 37521.7 | 11520.4 |
| JN.1.22 | Canada | 0.973 | 0.966 | 0.979 | 1.000 | 31075.5 | 12947.7 |
| JN.1.43.1 | Canada | 0.971 | 0.940 | 0.999 | 1.001 | 35622.5 | 12524.4 |
| XDD | Canada | 0.970 | 0.940 | 0.998 | 1.000 | 35348.9 | 11653.2 |
| JN.2.5 | Canada | 0.970 | 0.961 | 0.978 | 1.000 | 31010.2 | 13131.3 |
| JN.1.47 | Canada | 0.967 | 0.949 | 0.983 | 1.000 | 32464.2 | 11916.6 |
| JN.1.1 | Canada | 0.959 | 0.951 | 0.966 | 1.000 | 32263.5 | 12853.9 |
| JN.1.2 | Canada | 0.954 | 0.945 | 0.963 | 1.000 | 31722.5 | 11163.6 |
| EG.5.1.4 | Canada | 0.948 | 0.917 | 0.978 | 1.000 | 36896.3 | 12139.0 |
| JN.1.45 | Canada | 0.942 | 0.915 | 0.968 | 1.000 | 33895.1 | 10655.4 |
| JG.3 | Canada | 0.941 | 0.931 | 0.950 | 1.000 | 33874.1 | 12677.9 |
| BA.2.86.1 | Canada | 0.939 | 0.909 | 0.967 | 1.001 | 34430.6 | 11317.3 |
| JN.1.6 | Canada | 0.924 | 0.890 | 0.957 | 1.000 | 35939.9 | 12324.2 |
| EG.5.1.1 | Canada | 0.924 | 0.892 | 0.954 | 1.000 | 36641.2 | 11033.3 |
| JN.1.47.1 | Canada | 0.919 | 0.887 | 0.948 | 1.001 | 32113.8 | 12268.9 |
| HK.3.2 | Canada | 0.908 | 0.851 | 0.957 | 1.001 | 34426.8 | 11294.5 |
| EG.5.1.8 | Canada | 0.886 | 0.835 | 0.933 | 1.000 | 33891.0 | 11363.7 |
| HV.1 | Canada | 0.855 | 0.842 | 0.868 | 1.000 | 31478.6 | 12311.4 |
| JD.1.1 | Canada | 0.848 | 0.806 | 0.887 | 1.001 | 33912.1 | 11595.1 |
| JN.2 | Canada | 0.848 | 0.807 | 0.886 | 1.000 | 33276.3 | 12031.6 |
| HK.3 | Canada | 0.843 | 0.811 | 0.875 | 1.000 | 35315.1 | 11679.6 |
| FL.1.5.2 | Canada | 0.792 | 0.723 | 0.856 | 1.000 | 35291.0 | 11836.6 |
| JD.1.1.1 | Canada | 0.728 | 0.664 | 0.789 | 1.000 | 31028.3 | 11190.6 |
| JF.1 | Canada | 0.595 | 0.464 | 0.720 | 1.000 | 34182.7 | 11147.9 |
| LB.1 | United Kingdom | 1.439 | 1.348 | 1.547 | 1.001 | 21236.9 | 11393.9 |
| KP.2.3 | United Kingdom | 1.413 | 1.321 | 1.523 | 1.000 | 20850.3 | 12590.2 |
| KP.2.2 | United Kingdom | 1.400 | 1.294 | 1.531 | 1.000 | 20145.2 | 10476.8 |
| XDK.3 | United Kingdom | 1.375 | 1.274 | 1.503 | 1.000 | 23994.5 | 11279.9 |
| KP.3 | United Kingdom | 1.282 | 1.255 | 1.310 | 1.000 | 19716.2 | 11871.3 |
| XDK.1 | United Kingdom | 1.272 | 1.232 | 1.318 | 1.000 | 22067.0 | 11491.3 |
| KW.1.1 | United Kingdom | 1.268 | 1.199 | 1.352 | 1.000 | 23168.4 | 12144.1 |
| KP.4.1 | United Kingdom | 1.258 | 1.201 | 1.327 | 1.000 | 22914.5 | 9987.4 |
| KS.1 | United Kingdom | 1.246 | 1.213 | 1.283 | 1.000 | 18274.3 | 12009.4 |
| KP.1.1.1 | United Kingdom | 1.240 | 1.196 | 1.289 | 1.000 | 21198.9 | 11479.6 |
| JN.1.16.1 | United Kingdom | 1.230 | 1.209 | 1.252 | 1.000 | 18916.3 | 11815.2 |
| KP.1.1 | United Kingdom | 1.222 | 1.192 | 1.255 | 1.000 | 19739.6 | 10860.0 |
| LA.1 | United Kingdom | 1.211 | 1.167 | 1.262 | 1.000 | 19414.7 | 11320.7 |
| KP.2 | United Kingdom | 1.208 | 1.193 | 1.224 | 1.000 | 16573.6 | 11788.1 |
| JN.1.7.4 | United Kingdom | 1.206 | 1.162 | 1.258 | 1.000 | 20580.2 | 12031.9 |
| JN.1.13.1 | United Kingdom | 1.177 | 1.140 | 1.219 | 1.000 | 20329.2 | 11017.0 |
| JN.1.16 | United Kingdom | 1.133 | 1.119 | 1.147 | 1.000 | 17214.5 | 11762.0 |
| JN.1.7.1 | United Kingdom | 1.129 | 1.114 | 1.146 | 1.000 | 16967.2 | 12551.1 |

| | | | | | | | |
|---|---|---|---|---|---|---|---|
| JN.1.9.1 | United Kingdom | 1.122 | 1.101 | 1.143 | 1.000 | 20432.6 | 11158.2 |
| JN.1.11.1 | United Kingdom | 1.118 | 1.101 | 1.136 | 1.000 | 21142.6 | 12065.3 |
| JN.1.4.3 | United Kingdom | 1.110 | 1.080 | 1.142 | 1.000 | 21488.4 | 11677.3 |
| JN.1.20 | United Kingdom | 1.107 | 1.084 | 1.133 | 1.000 | 20986.8 | 12294.6 |
| JN.1.7 | United Kingdom | 1.106 | 1.099 | 1.113 | 1.000 | 12862.4 | 12340.4 |
| JN.1.34 | United Kingdom | 1.103 | 1.081 | 1.126 | 1.000 | 24045.7 | 11567.0 |
| JN.1.4.6 | United Kingdom | 1.098 | 1.084 | 1.112 | 1.000 | 18306.3 | 12185.1 |
| JN.1.4.4 | United Kingdom | 1.094 | 1.079 | 1.110 | 1.000 | 19464.7 | 11965.0 |
| JN.1.32.1 | United Kingdom | 1.086 | 1.075 | 1.097 | 1.000 | 16778.4 | 11885.6 |
| XDK | United Kingdom | 1.084 | 1.072 | 1.097 | 1.001 | 19434.1 | 12325.2 |
| JN.1.7.2 | United Kingdom | 1.081 | 1.061 | 1.102 | 1.000 | 22727.7 | 12451.8 |
| JN.1.18 | United Kingdom | 1.075 | 1.066 | 1.085 | 1.000 | 17825.0 | 12007.9 |
| JN.1.32 | United Kingdom | 1.061 | 1.053 | 1.070 | 1.000 | 15999.2 | 12414.3 |
| JN.1.6.1 | United Kingdom | 1.061 | 1.041 | 1.081 | 1.000 | 20094.7 | 12370.5 |
| JN.1.8.1 | United Kingdom | 1.059 | 1.042 | 1.076 | 1.000 | 23321.5 | 12013.6 |
| JN.1.5 | United Kingdom | 1.056 | 1.036 | 1.074 | 1.000 | 20618.8 | 11807.5 |
| JN.1.9 | United Kingdom | 1.044 | 1.032 | 1.056 | 1.001 | 18824.9 | 12916.2 |
| JN.1.45 | United Kingdom | 1.039 | 1.008 | 1.068 | 1.000 | 23960.1 | 13490.0 |
| XDS | United Kingdom | 1.039 | 1.016 | 1.061 | 1.000 | 22530.7 | 12643.3 |
| JN.1.39 | United Kingdom | 1.030 | 1.019 | 1.041 | 1.000 | 22032.7 | 11965.6 |
| JN.1.37 | United Kingdom | 1.024 | 0.995 | 1.051 | 1.000 | 23882.3 | 13365.7 |
| JN.1.4.5 | United Kingdom | 1.020 | 1.007 | 1.033 | 1.000 | 21439.3 | 13505.3 |
| JN.1.4 | United Kingdom | 1.017 | 1.011 | 1.023 | 1.000 | 15723.4 | 13069.9 |
| JN.1.1.1 | United Kingdom | 1.016 | 0.992 | 1.040 | 1.000 | 24389.4 | 12316.0 |
| JN.1.19 | United Kingdom | 1.015 | 0.999 | 1.030 | 1.000 | 23170.9 | 13205.9 |
| JN.1.8 | United Kingdom | 1.010 | 0.993 | 1.027 | 1.000 | 24155.4 | 12579.0 |
| XDT | United Kingdom | 1.008 | 0.972 | 1.041 | 1.000 | 23066.8 | 12045.2 |
| JN.1.4.1 | United Kingdom | 1.006 | 0.972 | 1.038 | 1.000 | 21522.3 | 12881.0 |
| JN.1.31 | United Kingdom | 1.001 | 0.970 | 1.030 | 1.000 | 21936.4 | 12171.0 |
| JN.1.22 | United Kingdom | 0.999 | 0.983 | 1.013 | 1.000 | 22265.5 | 12728.0 |
| JN.1.4.7 | United Kingdom | 0.998 | 0.972 | 1.022 | 1.000 | 22959.7 | 12852.0 |
| JN.1.43 | United Kingdom | 0.996 | 0.968 | 1.022 | 1.000 | 22885.8 | 12920.9 |
| JN.1.49 | United Kingdom | 0.994 | 0.964 | 1.020 | 1.000 | 23477.6 | 12850.8 |
| JN.1.2 | United Kingdom | 0.988 | 0.959 | 1.015 | 1.000 | 22728.0 | 12538.2 |
| JN.1.29 | United Kingdom | 0.979 | 0.935 | 1.018 | 1.000 | 20823.9 | 13121.8 |
| JN.1.8.3 | United Kingdom | 0.977 | 0.945 | 1.006 | 1.000 | 24591.9 | 12246.2 |
| XDD | United Kingdom | 0.970 | 0.930 | 1.006 | 1.000 | 21892.8 | 12574.5 |
| JN.4 | United Kingdom | 0.969 | 0.931 | 1.003 | 1.000 | 24001.2 | 12285.1 |
| XDN | United Kingdom | 0.960 | 0.940 | 0.980 | 1.000 | 23404.5 | 12405.8 |
| JN.1.1 | United Kingdom | 0.958 | 0.946 | 0.970 | 1.000 | 22717.1 | 13297.0 |
| BA.2.86.1 | United Kingdom | 0.957 | 0.933 | 0.980 | 1.000 | 23770.2 | 12999.1 |
| JN.2 | United Kingdom | 0.953 | 0.931 | 0.974 | 1.000 | 25814.6 | 12356.2 |
| JN.6 | United Kingdom | 0.950 | 0.920 | 0.978 | 1.000 | 23359.2 | 12761.0 |
| JN.3 | United Kingdom | 0.938 | 0.904 | 0.970 | 1.000 | 25774.4 | 11832.0 |
| JN.10 | United Kingdom | 0.919 | 0.862 | 0.968 | 1.001 | 22371.1 | 12357.7 |
| HV.1 | United Kingdom | 0.906 | 0.862 | 0.948 | 1.000 | 23045.9 | 13523.9 |
| JG.3 | United Kingdom | 0.905 | 0.864 | 0.943 | 1.000 | 19396.7 | 12574.3 |
| JN.18 | United Kingdom | 0.878 | 0.822 | 0.930 | 1.000 | 21076.7 | 12471.9 |
| JD.1.1 | United Kingdom | 0.878 | 0.829 | 0.923 | 1.000 | 22371.4 | 12778.2 |
| XBB.1.16.17 | United Kingdom | 0.826 | 0.750 | 0.896 | 1.000 | 21391.5 | 11874.2 |
| KP.3.2 | USA | 1.650 | 1.487 | 1.848 | 1.000 | 16811.5 | 10427.5 |
| LB.1 | USA | 1.331 | 1.297 | 1.368 | 1.000 | 14361.3 | 11867.3 |
| KP.2.3 | USA | 1.306 | 1.266 | 1.351 | 1.000 | 16292.9 | 11993.4 |
| XDV.1 | USA | 1.297 | 1.244 | 1.356 | 1.000 | 16402.7 | 11451.3 |
| KP.4.1 | USA | 1.258 | 1.204 | 1.321 | 1.000 | 15406.4 | 10956.6 |
| KP.2.2 | USA | 1.247 | 1.212 | 1.286 | 1.000 | 15008.6 | 12383.0 |
| KW.1.1 | USA | 1.227 | 1.176 | 1.285 | 1.000 | 17584.4 | 11162.6 |
| KP.1.1.1 | USA | 1.224 | 1.191 | 1.259 | 1.000 | 15832.4 | 12138.3 |
| KP.3 | USA | 1.208 | 1.187 | 1.229 | 1.000 | 14213.1 | 11551.4 |
| KS.1 | USA | 1.204 | 1.180 | 1.229 | 1.000 | 15111.7 | 11910.3 |
| LA.1 | USA | 1.203 | 1.160 | 1.253 | 1.000 | 15563.3 | 11177.8 |
| JN.1.16.1 | USA | 1.182 | 1.166 | 1.199 | 1.000 | 16024.2 | 12572.0 |
| KP.2 | USA | 1.175 | 1.166 | 1.185 | 1.000 | 13573.7 | 10323.3 |
| KP.1.1 | USA | 1.173 | 1.156 | 1.190 | 1.000 | 15817.4 | 12026.0 |
| JN.1.1.6 | USA | 1.170 | 1.143 | 1.198 | 1.000 | 15789.6 | 11914.7 |
| JN.1.40 | USA | 1.134 | 1.108 | 1.163 | 1.000 | 15240.0 | 11646.2 |
| JN.1.16 | USA | 1.133 | 1.124 | 1.143 | 1.000 | 13835.7 | 10886.3 |
| JN.1.18.2 | USA | 1.133 | 1.104 | 1.164 | 1.000 | 17703.9 | 12831.7 |
| JN.1.11.1 | USA | 1.111 | 1.100 | 1.123 | 1.000 | 16906.7 | 12338.5 |
| KW.1 | USA | 1.108 | 1.086 | 1.129 | 1.000 | 14059.8 | 11012.2 |
| JN.1.4.4 | USA | 1.103 | 1.096 | 1.110 | 1.000 | 14947.0 | 12026.6 |
| JN.1.11 | USA | 1.101 | 1.083 | 1.119 | 1.000 | 14462.0 | 11963.3 |
| JN.1.13.1 | USA | 1.083 | 1.078 | 1.088 | 1.000 | 13678.4 | 11585.2 |
| JN.1.4.3 | USA | 1.079 | 1.069 | 1.089 | 1.001 | 16632.8 | 12107.2 |
| JN.1.24.1 | USA | 1.076 | 1.055 | 1.097 | 1.000 | 19399.9 | 12749.1 |
| KP.1 | USA | 1.075 | 1.062 | 1.088 | 1.000 | 15675.1 | 11968.6 |
| JN.1.7.2 | USA | 1.074 | 1.064 | 1.084 | 1.001 | 16814.3 | 13512.0 |
| JN.1.7 | USA | 1.074 | 1.070 | 1.077 | 1.000 | 10780.6 | 12268.6 |
| JN.1.32 | USA | 1.072 | 1.065 | 1.079 | 1.000 | 15721.3 | 13190.2 |
| JN.1.18 | USA | 1.069 | 1.063 | 1.076 | 1.000 | 13987.2 | 12172.7 |
| JN.1.7.3 | USA | 1.069 | 1.039 | 1.099 | 1.001 | 16127.4 | 11978.8 |
| XDK | USA | 1.066 | 1.049 | 1.081 | 1.000 | 18228.7 | 12783.9 |
| XDP.1 | USA | 1.064 | 1.050 | 1.077 | 1.001 | 15981.3 | 12049.5 |
| XDR | USA | 1.062 | 1.032 | 1.091 | 1.001 | 18234.2 | 12703.3 |
| JN.1.37 | USA | 1.060 | 1.041 | 1.078 | 1.000 | 18152.6 | 12885.0 |
| JN.1.4.6 | USA | 1.060 | 1.048 | 1.071 | 1.000 | 13748.3 | 12761.4 |
| JN.1.32.1 | USA | 1.059 | 1.031 | 1.086 | 1.000 | 17470.2 | 12026.1 |
| XDQ | USA | 1.058 | 1.042 | 1.075 | 1.000 | 17009.3 | 11951.8 |
| KR.1 | USA | 1.056 | 1.026 | 1.086 | 1.000 | 15838.7 | 12341.5 |
| JN.1.13 | USA | 1.051 | 1.025 | 1.077 | 1.000 | 16613.2 | 12659.0 |
| KV.2 | USA | 1.048 | 1.042 | 1.055 | 1.000 | 15300.4 | 11201.2 |
| JN.1.15 | USA | 1.048 | 1.030 | 1.064 | 1.000 | 17152.0 | 12779.2 |
| JN.1.8.1 | USA | 1.047 | 1.043 | 1.051 | 1.000 | 13214.0 | 11962.2 |

| | | | | | | | |
|---|---|---|---|---|---|---|---|
| JN.1.30 | USA | 1.046 | 1.024 | 1.068 | 1.000 | 16472.8 | 11199.2 |
| JN.1.14 | USA | 1.046 | 1.030 | 1.061 | 1.000 | 18692.3 | 12502.8 |
| JN.1.20 | USA | 1.042 | 1.028 | 1.056 | 1.000 | 17095.0 | 12248.6 |
| JN.1.34 | USA | 1.038 | 1.024 | 1.053 | 1.000 | 17347.9 | 12080.6 |
| JN.1.26 | USA | 1.038 | 1.018 | 1.058 | 1.000 | 16421.2 | 12343.3 |
| JN.1.8.3 | USA | 1.036 | 1.012 | 1.059 | 1.000 | 17365.7 | 12655.6 |
| JN.1.1.5 | USA | 1.035 | 1.018 | 1.051 | 1.000 | 15866.7 | 12267.7 |
| JN.1.10 | USA | 1.026 | 0.996 | 1.054 | 1.000 | 17335.0 | 12287.2 |
| XDP | USA | 1.023 | 1.015 | 1.031 | 1.000 | 16682.7 | 12447.7 |
| BA.2 | USA | 1.023 | 1.007 | 1.039 | 1.000 | 17314.1 | 12328.7 |
| KQ.1 | USA | 1.020 | 1.006 | 1.033 | 1.000 | 15320.0 | 12616.7 |
| JN.1.43.1 | USA | 1.019 | 1.008 | 1.030 | 1.000 | 17945.7 | 12645.7 |
| JN.1.4.2 | USA | 1.019 | 1.013 | 1.025 | 1.000 | 15728.3 | 12411.7 |
| JN.1.8 | USA | 1.017 | 1.009 | 1.026 | 1.000 | 16160.5 | 12112.7 |
| JN.1.5 | USA | 1.016 | 1.004 | 1.028 | 1.000 | 16526.6 | 12917.9 |
| JN.1.4.7 | USA | 1.015 | 1.001 | 1.029 | 1.000 | 17764.8 | 12753.0 |
| JN.1.28 | USA | 1.013 | 0.981 | 1.044 | 1.000 | 15804.3 | 13322.8 |
| JN.1.6.1 | USA | 1.012 | 0.981 | 1.041 | 1.000 | 17585.6 | 12202.6 |
| JN.1.39 | USA | 1.010 | 1.004 | 1.015 | 1.000 | 15317.9 | 12853.5 |
| JN.1.42 | USA | 1.010 | 1.003 | 1.016 | 1.000 | 15458.1 | 12464.0 |
| JN.1.21 | USA | 1.009 | 0.990 | 1.028 | 1.000 | 17539.5 | 12771.4 |
| JN.1.29 | USA | 1.009 | 0.992 | 1.025 | 1.000 | 16106.1 | 12647.8 |
| JN.1.3 | USA | 1.009 | 0.990 | 1.026 | 1.000 | 16572.2 | 12790.3 |
| GE.1.2 | USA | 1.006 | 0.978 | 1.033 | 1.000 | 16967.8 | 12575.8 |
| JN.1.4 | USA | 1.003 | 1.000 | 1.005 | 1.000 | 10222.3 | 12037.2 |
| JN.1.27 | USA | 1.001 | 0.979 | 1.023 | 1.000 | 17635.3 | 13053.7 |
| JN.1.6 | USA | 1.001 | 0.992 | 1.009 | 1.000 | 15671.9 | 12852.8 |
| JN.1.4.5 | USA | 0.999 | 0.995 | 1.003 | 1.000 | 13876.7 | 12402.5 |
| JN.1.9 | USA | 0.997 | 0.991 | 1.004 | 1.000 | 17307.1 | 13453.6 |
| JN.1.17 | USA | 0.997 | 0.983 | 1.010 | 1.000 | 17797.9 | 12767.6 |
| JN.1.4.1 | USA | 0.996 | 0.970 | 1.020 | 1.000 | 17726.1 | 12545.5 |
| JN.1.22 | USA | 0.995 | 0.986 | 1.003 | 1.001 | 18033.7 | 12349.0 |
| JN.1.19 | USA | 0.994 | 0.983 | 1.004 | 1.001 | 15127.9 | 12829.1 |
| XDD | USA | 0.993 | 0.979 | 1.007 | 1.000 | 18056.0 | 12394.5 |
| JN.1.46 | USA | 0.992 | 0.983 | 1.001 | 1.001 | 17013.4 | 12471.5 |
| JN.1.1.1 | USA | 0.991 | 0.961 | 1.020 | 1.000 | 16333.9 | 12527.7 |
| JN.1.31 | USA | 0.988 | 0.976 | 1.000 | 1.000 | 17995.9 | 13175.3 |
| JN.1.38 | USA | 0.987 | 0.977 | 0.996 | 1.000 | 17716.1 | 12996.8 |
| JN.1.42.1 | USA | 0.985 | 0.944 | 1.021 | 1.000 | 17149.2 | 12215.3 |
| JN.1.43 | USA | 0.981 | 0.972 | 0.989 | 1.001 | 17685.1 | 12389.7 |
| JN.1.49 | USA | 0.980 | 0.949 | 1.009 | 1.001 | 17461.5 | 12140.8 |
| JN.4 | USA | 0.980 | 0.939 | 1.017 | 1.000 | 16659.3 | 13543.0 |
| JN.2.5 | USA | 0.978 | 0.946 | 1.008 | 1.001 | 19387.4 | 13534.5 |
| JN.1.2 | USA | 0.977 | 0.969 | 0.985 | 1.000 | 15455.0 | 12683.6 |
| JN.1.45 | USA | 0.976 | 0.962 | 0.989 | 1.000 | 19216.8 | 12686.4 |
| JN.1.47.2 | USA | 0.973 | 0.935 | 1.008 | 1.000 | 15482.2 | 11879.6 |
| JN.1.47 | USA | 0.969 | 0.955 | 0.982 | 1.000 | 18003.9 | 12317.4 |
| JN.1.24 | USA | 0.967 | 0.936 | 0.996 | 1.000 | 16845.4 | 12868.9 |
| XBB.1.41.1 | USA | 0.961 | 0.936 | 0.986 | 1.000 | 18381.4 | 13643.9 |
| BA.2.86.1 | USA | 0.958 | 0.942 | 0.973 | 1.000 | 17483.6 | 12830.6 |
| JN.1.1 | USA | 0.958 | 0.953 | 0.963 | 1.000 | 15770.9 | 13319.8 |
| JN.1.41 | USA | 0.947 | 0.929 | 0.964 | 1.000 | 18358.1 | 12803.0 |
| JN.3 | USA | 0.934 | 0.913 | 0.954 | 1.000 | 18384.2 | 13380.3 |
| EG.5.1.3 | USA | 0.929 | 0.879 | 0.975 | 1.000 | 20190.9 | 12583.0 |
| HK.3.1 | USA | 0.925 | 0.872 | 0.972 | 1.000 | 16061.6 | 13048.3 |
| JN.2 | USA | 0.923 | 0.894 | 0.950 | 1.000 | 19272.1 | 13875.0 |
| JN.1.1.7 | USA | 0.922 | 0.866 | 0.973 | 1.000 | 17815.3 | 12654.7 |
| EG.5.1.6 | USA | 0.921 | 0.889 | 0.952 | 1.000 | 18963.2 | 13327.2 |
| FY.5 | USA | 0.920 | 0.870 | 0.965 | 1.000 | 18255.3 | 12410.8 |
| JG.3 | USA | 0.911 | 0.899 | 0.922 | 1.000 | 18132.6 | 13813.0 |
| JE.1.1 | USA | 0.906 | 0.847 | 0.960 | 1.000 | 19266.4 | 11983.9 |
| GK.1.1 | USA | 0.905 | 0.869 | 0.938 | 1.000 | 17846.0 | 13128.8 |
| GK.1 | USA | 0.896 | 0.836 | 0.949 | 1.000 | 17908.4 | 12786.7 |
| XBB.1.16.6 | USA | 0.895 | 0.866 | 0.922 | 1.000 | 19713.5 | 13270.3 |
| EG.5.1 | USA | 0.893 | 0.862 | 0.922 | 1.000 | 18579.7 | 12278.1 |
| HK.1 | USA | 0.892 | 0.830 | 0.946 | 1.001 | 16287.9 | 11275.4 |
| HK.3.2 | USA | 0.883 | 0.850 | 0.914 | 1.000 | 16341.1 | 12341.0 |
| XBB.1.16.17 | USA | 0.880 | 0.840 | 0.919 | 1.000 | 19290.7 | 11523.6 |
| JD.1.1 | USA | 0.880 | 0.864 | 0.896 | 1.000 | 19958.3 | 13322.9 |
| XBB.1.16.11 | USA | 0.880 | 0.839 | 0.919 | 1.000 | 18596.0 | 12117.8 |
| FL.1.5.1 | USA | 0.879 | 0.849 | 0.907 | 1.000 | 19285.5 | 12503.1 |
| FL.1.5.2 | USA | 0.878 | 0.832 | 0.921 | 1.000 | 18541.9 | 13155.2 |
| JF.1 | USA | 0.874 | 0.844 | 0.903 | 1.000 | 20321.6 | 12820.1 |
| EG.5.1.1 | USA | 0.873 | 0.845 | 0.901 | 1.000 | 18073.2 | 13704.2 |
| JN.18 | USA | 0.872 | 0.813 | 0.926 | 1.001 | 20610.4 | 12012.8 |
| HV.1 | USA | 0.872 | 0.864 | 0.879 | 1.000 | 17697.4 | 13899.1 |
| HK.3 | USA | 0.871 | 0.852 | 0.890 | 1.000 | 17651.8 | 12728.8 |
| JD.1.1.1 | USA | 0.871 | 0.842 | 0.898 | 1.000 | 17611.4 | 12150.7 |
| XBB.1.16.15 | USA | 0.864 | 0.813 | 0.910 | 1.000 | 19758.2 | 13020.8 |
| EG.5.1.8 | USA | 0.829 | 0.767 | 0.885 | 1.000 | 18437.1 | 11830.9 |
| FL.15.1.1 | USA | 0.785 | 0.708 | 0.855 | 1.000 | 17706.5 | 12159.4 |

The relative $R_e$ of JN.1 is set to 1.

**Table S4. Primers used in this study**

| Primer name | Primer sequence (5'-to-3') | Purpose |
|---|---|---|
| Omicron universal Fw | cactatagggcgaattgggtaccatgtttgtgttcctggt | Preparation of S expression plasmid |
| BA.2 WT Rv | agctccaccgcggtggcggccgctcaggtgtagtgcagtttca | Preparation of S expression plasmid |
| KP2_Q493E Fw | tgttactttccactcGAGtcctatGGCttcaga | Preparation of S expression plasmid |
| KP2_Q493E Rv | tctgaaGCCataggaCTCgagtggaaagtaaca | Preparation of S expression plasmid |
| KP2_delS31_Fw | caaAGCtacaccaacttcaccaggggagtc | Preparation of S expression plasmid |
| KP2_delS31_Rv | gactcccctggtgaagttggtgtaGCTttg | Preparation of S expression plasmid |
| KP2_H146Q_Fw | ttcctgGACgtctacCAGaagaacaacaagtcc | Preparation of S expression plasmid |
| KP2_H146Q_Rv | ggacttgttgttcttCTGgtagacGTCcaggaa | Preparation of S expression plasmid |
| JN1_Q183H_Fw | gacttggagggcaagCACggcaacttcaagaac | Preparation of S expression plasmid |
| JN1_Q183H_Rv | gttcttgaagttgccGTGcttgccctccaagtc | Preparation of S expression plasmid |

### Consortia

#### The Genotype to Phenotype Japan (G2P-Japan) Consortium

**The Institute of Medical Science, The University of Tokyo, Japan**
Naoko Misawa, Arnon Plianchaisuk, Ziyi Guo, Alfredo Hinay Jr., Kaoru Usui, Wilaiporn Saikruang, Spyridon Lytras, Daichi Yamasoba, Yusuke Kosugi, Shusuke Kawakubo, Luca Nishimura, Shigeru Fujita, Luo Chen, Lin Pan, Wenye Li, Kio Horinaka, Mai Suganami, Mika Chiba, Kyoko Yasuda, Keiko Iida, Adam P. Strange, Naomi Ohsumi, Shiho Tanaka, Eiko Ogawa, Tsuki Fukuda, Rina Osujo

**Hokkaido University, Japan**
Takasuke Fukuhara, Tomokazu Tamura, Rigel Suzuki, Saori Suzuki, Shuhei Tsujino, Hayato Ito, Hirofumi Sawa, Naganori Nao, Keita Matsuno, Keita Mizuma, Jingshu Li, Izumi Kida, Yume Mimura, Yuma Ohari, Shinya Tanaka, Masumi Tsuda, Lei Wang, Yoshikata Oda, Zannatul Ferdous, Kenji Shishido, Hiromi Mohri, Miki Iida

**Tokyo Metropolitan Institute of Public Health**
Kenji Sadamasu, Kazuhisa Yoshimura, Hiroyuki Asakura, Mami Nagashima, Isao Yoshida

**Tokai University, Japan**
So Nakagawa

**Kyoto University, Japan**
Kotaro Shirakawa, Akifumi Takaori-Kondo, Kazuo Takayama, Rina Hashimoto, Sayaka Deguchi, Yukio Watanabe, Yoshitaka Nakata, Hiroki Futatsusako, Ayaka Sakamoto, Naoko Yasuhara, Takao Hashiguchi, Tateki Suzuki, Kanako Kimura, Jiei Sasaki, Yukari Nakajima, Hisano Yajima

**Hiroshima University, Japan**
Takashi Irie, Ryoko Kawabata

**Kyushu University, Japan**
Kaori Tabata

**Kumamoto University, Japan**
Terumasa Ikeda, Hesham Nasser, Ryo Shimizu, MST Monira Begum, Michael Jonathan, Yuka Mugita, Sharee Leong, Otowa Takahashi, Takamasa Ueno, Chihiro Motozono, Mako Toyoda

**University of Miyazaki, Japan**
Akatsuki Saito, Anon Kosaka, Miki Kawano, Natsumi Matsubara, Tomoko Nishiuchi

**Charles University, Czechia**
Jiri Zahradnik, Prokopios Andrikopoulos, Miguel Padilla-Blanco, Aditi Konar, Ruojin Tuan

**Acknowledgments**

We would like to thank all members of The Genotype to Phenotype Japan (G2P-Japan) Consortium. We thank Kenzo Tokunaga (National Institute of Infectious Diseases, Japan) for sharing materials and Yuka Kamoshita (Department of Laboratory Medicine, Keio University School of Medicine) and Masayo Noguchi (Clinical Laboratory, Keio University Hospital) for supporting patient sera collection. We gratefully acknowledge the numerous laboratories worldwide that have provided sequence data and metadata to GISAID. A full list of originating and submitting laboratories for the sequences used in our analysis can be found at https://www.gisaid.org using the EPI-SET-ID: EPI_SET_240529sa.

**Supplementary References**


1. Suzuki R, Yamasoba D, Kimura I, et al. Attenuated fusogenicity and pathogenicity of SARS-CoV-2 Omicron variant. *Nature* 2022; **603**(7902): 700-5.
2. Saito A, Irie T, Suzuki R, et al. Enhanced fusogenicity and pathogenicity of SARS-CoV-2 Delta P681R mutation. *Nature* 2022; **602**(7896): 300-6.
3. Yamasoba D, Kimura I, Nasser H, et al. Virological characteristics of the SARS-CoV-2 Omicron BA.2 spike. *Cell* 2022; **185**(12): 2103-15.e19.
4. Kimura I, Yamasoba D, Tamura T, et al. Virological characteristics of the SARS-CoV-2 Omicron BA.2 subvariants, including BA.4 and BA.5. *Cell* 2022; **185**(21): 3992-4007.e16.
5. Uriu K, Ito J, Kosugi Y, et al. Transmissibility, infectivity, and immune evasion of the SARS-CoV-2 BA.2.86 variant. *Lancet Infect Dis* 2023; **23**(11): e460-e1.
6. Kosugi Y, Plianchaisuk A, Putri O, et al. Characteristics of the SARS-CoV-2 omicron HK.3 variant harbouring the FLip substitution. *Lancet Microbe* 2024.
7. Kaku Y, Kosugi Y, Uriu K, et al. Antiviral efficacy of the SARS-CoV-2 XBB breakthrough infection sera against omicron subvariants including EG.5. *Lancet Infect Dis* 2023; **23**(10): e395-e6.
8. Kaku Y, Okumura K, Padilla-Blanco M, et al. Virological characteristics of the SARS-CoV-2 JN.1 variant. *Lancet Infect Dis* 2024; **24**(2): e82.
9. Niwa H, Yamamura K, Miyazaki J. Efficient selection for high-expression transfectants with a novel eukaryotic vector. *Gene* 1991; **108**(2): 193-9.
10. Ozono S, Zhang Y, Ode H, et al. SARS-CoV-2 D614G spike mutation increases entry efficiency with enhanced ACE2-binding affinity. *Nat Commun* 2021; **12**(1): 848.
11. Ferreira I, Kemp SA, Datir R, et al. SARS-CoV-2 B.1.617 Mutations L452R and E484Q Are Not Synergistic for Antibody Evasion. *J Infect Dis* 2021; **224**(6): 989-94.
12. Motozono C, Toyoda M, Zahradnik J, et al. SARS-CoV-2 spike L452R variant evades cellular immunity and increases infectivity. *Cell Host Microbe* 2021; **29**(7): 1124-36.e11.
13. Ozono S, Zhang Y, Tobiume M, Kishigami S, Tokunaga K. Super-rapid quantitation of the production of HIV-1 harboring a luminescent peptide tag. *J Biol Chem* 2020; **295**(37): 13023-30.
14. Garcia-Beltran WF, St Denis KJ, Hoelzemer A, et al. mRNA-based COVID-19 vaccine boosters induce neutralizing immunity against SARS-CoV-2 Omicron variant. *Cell* 2022; **185**(3): 457-66.e4.
15. Kaku Y, Uriu K, Kosugi Y, et al. Virological characteristics of the SARS-CoV-2 KP.2 variant. *Lancet Infect Dis* 2024.
